# Applying biostimulants boosts forage productivity without affecting soil biotic and abiotic parameters on a Central Coast California rangeland

**DOI:** 10.1101/2021.12.31.474676

**Authors:** Chelsea J. Carey, Hayley Strohm, Ford Smith, Mark Biaggi

## Abstract

There is increasing interest in using biostimulant products, such as microbial inoculants and humic substances, to help manage rangelands regeneratively. Understanding how plant and soil communities on rangelands respond to these products is therefore important. In this study, we examined the combined effects of a commercial inoculant and humic product that are currently on the market, and asked whether they influenced rangeland forage productivity and quality, soil microbial biomass and community composition, and abiotic soil parameters in Central Coastal California. We found that forage productivity and some metrics of forage quality responded positively to the foliar application of a commercial microbial inoculant and humic product, but that these benefits were not mirrored by changes belowground in the microbial community or abiotic parameters. Depending on the goals of using the products, this could be seen as a winning scenario and suggests microbial inoculants and humic products could warrant attention as a potential tool for regenerative stewardship of rangelands. While our study derives from one ranch and therefore requires confirmation of its ubiquity prior to broadscale adoption, our results provide new insights into the usefulness of this approach for managing rangeland productivity in California’s Central Coast.

## Introduction

Regenerative agriculture aims to restore soil health and biodiversity, sequester carbon from the atmosphere, maximize water and nutrient use efficiency, and provide ample and nutritious food while reducing reliance on inputs from chemical fertilizers and pesticides. Management strategies used by those practicing regenerative agriculture are as varied as the production systems themselves, but tend to include practices that minimize soil disturbance, maximize crop diversity, keep the soil covered, maintain living roots, and integrate livestock. The use of microbial inoculants and humic products (derived from alkali extractions of soil humic acid [HA] and fulvic acid [FA] fractions) are receiving increased attention as a potential tool for regenerative management both of croplands and rangelands, because of their expected ability to act as biostimulants and enhance yield, crop quality, and soil conditions while simultaneously reducing the need for chemical inputs (Adesemoye et al. 2009; Calvo et al. 2014; Arora et al. 2020).

These expectations are supported in part by decades of research on the use of microbial inoculants and humic products to improve crop productivity and quality (reviewed in Hart & Forsythe 2012; Kong et al. 2018; Jindo et al. 2020). For example, research has shown that inoculants have the potential to improve nutrient availability and plant uptake through the addition of specific microbial taxa with, for example, capabilities to fix atmospheric nitrogen (N2), solubilize phosphorus, or stimulate root growth and elongation (Hamdali et al. 2008; Halpern et al. 2015; Chamizo et al. 2018). Inoculation with putative plant growth promoting (PGP) microorganisms can also alleviate plant environmental stress and prevent infections by phytopathogens through the synthesis of targeted enzymes, signaling molecules, and other compounds such as siderophores (reviewed in Khatoon et al. 2020; Arora et al. 2020).

Like microbial inoculants, humic products have been shown to improve crop productivity and quality (Rose et al. 2014) through a number of proposed mechanisms, which have been reviewed extensively elsewhere (Jindo et al. 2020). These mechanisms include improved nutrient availability, particularly of iron, through increased solubility and mobility in the soil (Nardi et al. 2002; Halpern et al. 2015), as well as enhanced plant uptake of nutrients through root growth (Nardi et al. 2002; Shah et al. 2018). The ability of plants to withstand stressful conditions is also aided by humic products and their effects on secondary metabolism of specialized plant compounds such as auxins (Del Buono 2021).

Despite promising—albeit equivocal (Hartz and Bottoms 2010; Haider et al. 2014)—evidence from the literature, the reliability of commercial microbial inoculants and humic products to consistently achieve desired outcomes in agricultural settings remains in question (Olk et al. 2018; Hart et al. 2018). In part, this is because the efficacy of inoculants has been shown to depend on the source (i.e., commercial or homemade), content (i.e., the community members present), and concentration of the product (Maltz et al. 2015). Moreover, introducing novel microorganisms with inoculants may result in unintended ecological consequences, possibly negating any observed benefits (Hart et al. 2018; Kaminsky et al. 2019). The effects of humic products will also depend on the source, concentration, and molecular weight of the humic fraction—in addition to the mode of application (Jindo et al. 2020). To complicate the picture further, characteristics of the recipient ecosystem have been shown to mediate the outcome of product application as well (Nardi et al. 2002; Olk et al. 2018; Olk et al. 2021).

As the chemical and biological mechanisms for efficacy continue to be researched (Mao et al. 2013), field studies that aim to evaluate commercial inoculants and humic products under operational conditions can help to disentangle these complex interactions from an applied perspective (Olk et al. 2013; Mayer et al. 2010). This is particularly needed in rangelands, where the use of such products is nascent but rapidly emerging as ranchers, policy-makers, and scientists alike look for new and alternative management practices that help promote outcomes related to forage production, climate stability, and soil health in the face of drought and other environmental threats (Godde et al. 2020).

The goal of this study was to evaluate the combined effects of a commercially available inoculant and humic product on rangeland dynamics in Central Coastal California. While the detailed composition of these products is proprietary and thus not available to the public, the inoculant is described by the manufacturer as containing ‘a stable solution of extracted beneficial microbes’ that include *Bacillus subtilis, Mucor heimalis*, and *Trichoderma harzianum*. The humic product is described as a ‘water-soluble powder that is composed of humic acid derived from Leonardite, kelp, complex carbohydrates, and amino acids.’ The specific objectives of our study were to track the combined effects of these products on (1) forage productivity, (2) forage quality, (3) soil microbial parameters associated with biomass and community structure (composition and diversity), and (4) abiotic soil properties during three consecutive years of application.

## Methods

### Study Site & Experimental Design

This study was conducted at TomKat Ranch (https://tomkatranch.org/), a 728 hectare property located in Pescadero, California, USA (37.261428, −122.360451). TomKat Ranch runs a grass-fed/grass-finished cow-calf operation using a planned grazing approach (Henneman & Seavy 2014). The site experiences a maritime Mediterranean-type climate characterized by mild relatively dry summers and cool wet winters. Mean annual air temperature is 12.9 °C and mean annual precipitation is 750 mm. During the 2017-18, 2018-19, and 2019-20 water years (July - June), cumulative rainfall totalled 491 mm, 877 mm, and 366 mm, respectively (Appendix Figure 1). The ranch contains approximately 324 hectares of grassland dominated by exotic annual grasses (e.g., *Bromus spp.; Brachypodium spp; Avena spp*) with some less-abundant native and non-native perennial grass species (primarily *Stipa pulchra, Danthonia californica, Phalaris aquatica*) and forbs. There are 11 soil series encompassed within the ranch boundaries, all of which derive from sedimentary rock, including mudstone, sandstone, and shale. The dominant soil series are Colma, Pomponio, Diablo, and Santa Lucia, with soil texture predominantly clay loam (sand = 15-60% [median 28.8%], silt = 16-45% [median 36.2%], clay = 18-54% [median 33.8%]).

Treatment and control plots (0.4 hectare) were established in a paired design, with one control and one treatment plot in each of ten fields. Near the center of each 0.4-ha plot, a 10 m x10 m soil sampling area was delineated and, adjacent to the sampling area, a grazing exclosure was erected (0.8 m radius; Appendix Figure 2). Prior to each growing season, the grazing exclosures were moved to a new adjacent area to minimize any effects that may accrue from prolonged exclusion. Careful site selection ensured that ecosystem characteristics such as slope, aspect, and vegetation type remained consistent between the 10 m x 10 m sampling plots and exclosures for each field, but not necessarily across fields.

Between February 20 and 24, 2018, we established the treatments by applying Soil Provide^®^ (Earthfort Inc., Corvallis, CO, USA), a commercially available inoculant, and Soil Revive^®^ (Earthfort Inc., Corvallis, CO, USA), a commercially available humic product, either separately or in combination, to each treatment plot. The Revive mixture contained 6% humic acid derived from leonardite, and nutrients derived from ascophyllum nodosum and potassium hydroxide in the following concentrations: 0.5% total nitrogen, 0.5 % available phosphate (P_2_O_5_), and 4% soluble potash (K_2_O). The initial application rate in February 2018 varied across fields to mimic real-world recommendations made to farmers and ranchers by the manufacturer (Appendix Table 1). Subsequent to the initial application, we applied the humic acid and microbial inoculant to all ten treatment plots at a constant rate of 1.12 kg per hectare and 9.35 liters per hectare, respectively, two to three times during each growing season (Appendix Figure 3). Products were suspended and applied in 136 liters of water; however, given the intention of the experiment was to test overall treatment effects (which inherently includes water as a part of it), and given 136 L is a minimal amount compared to background levels of fog deposition (Hiatt et al. 2012), we did not add water to the controls. All treatments were applied using a spray tank with a constant spray rate and 3.7 m wide swath.

### Aboveground Plant Biomass & Forage Quality

#### Plant Biomass

We collected plant metrics near the time of peak biomass across three growing seasons to determine treatment effects (Appendix Figure 2). In May 2018, 2019, and 2020, we measured aboveground plant biomass. Briefly, vegetation clippings were taken to the soil surface from within a 16.5 cm diameter ring at the center of each grazing exclosure. The biomass samples were then air dried for 3 days and oven dried at 60 °C to constant mass before weighing. Due to covid-19 processing complications, we were unable to obtain reliable results for the 2020 plant biomass estimates and therefore excluded them from subsequent analysis.

#### Forage Quality

In May 2018, 2019, and 2020, we also assessed metrics of forage quality. To do so, four randomly-selected vegetation clippings per treatment were taken from just outside the soil sampling area, composited, and shipped to Ward Laboratories. Because they were randomly selected, clippings encompassed numerous and variable plant species. At Ward Laboratories, the samples were analyzed via near infrared reflectance spectrometry (NIRS; Weiss and Hall 2020). Metrics of forage quality included those related to digestibility (acid detergent fiber [ADF], neutral detergent fiber [NDF], lignin), protein, fat, and mineral content (crude protein, crude fat, potassium [K^+^], calcium [Ca^2+^], magnesium [Mg^2+^], and phosphorus [P]). Raw results are reported on a % dry matter basis.

### Microbial & Abiotic Soil Properties

We collected soils across the same three growing seasons to determine belowground treatment effects (Appendix Figure 2), although not all analyses were conducted at every time point. Specifically, we collected soils in early February 2018 (just prior to treatment application) and on five subsequent dates in May 2018, November 2018, May 2019, December 2019, and May 2020. Fall sampling dates were timed to occur after the first rains. Each treatment had ten replicates (n = 10), where one replicate was a composite of ten soil cores (1.9 cm diameter x 10 cm deep) taken from random locations within the 10 m x 10 m sampling area. This sampling strategy provided us with a good estimate of the mean soil properties of the sampling area, which was our unit of replication. Composited soil samples were mixed, stored, and shipped for analysis as described below.

#### Microbial Biomass

At each collection date, we subsampled and labelled soils using a treatment-anonymous labeling scheme, and shipped them to Earthfort Inc. (Corvallis, OR, USA) for analysis of active and total microbial biomass using direct enumeration, which allows for the separation quantification of bacteria, fungi, and protozoa. Soil subsamples were stored at 4 °C until shipment, which occurred within 24 hours of collection. For all sampling dates, total and active bacteria were estimated by direct enumeration using fluorescein isothiocyanate (FITC; Babiuk and Paul 1970) and fluorescein diacetate (FDA; Ingham and Klein 1984) methods, respectively. Total and active fungi were calculated by measuring the diameter and length of hyphae (Lodge and Ingham 1991), with the active fungi method also using FDA. In December of 2019 and May of 2020, subgroups of protozoa (flagellates, amoeba, ciliates) were also enumerated by direct counting of serial dilutions. The subgroups are based on body type and method of movement, and the direct counts were used to estimate population sizes for each subgroup using the most probable number approach (Darbyshire et al. 1974).

#### Microbial Community Structure

To determine treatment effects on soil bacterial/archaeal and fungal community structure, soil samples from February 2018, May 2018, and November 2018 were analyzed using DNA amplicon sequencing. Subsamples were frozen at −20 °C prior to being shipped overnight on dry ice to Argonne National Laboratory’s Environmental Sample Preparation and Sequencing Facility (Lemont, IL, USA). Here, microbial DNA was extracted and amplicon libraries prepared for the V4 region of the 16S rRNA gene (forward barcoded 515f-806r primer pair; Walters et al. 2016) and internal transcribed spacer (ITS1) region (reverse barcoded ITS1f-ITS2 primer pair; Smith & Peay 2014) following Earth Microbiome Project protocols (Gilbert et al. 2010). Amplicons were subsequently sequenced on an Illumina MiSeq PE 2×150 run (16S rRNA) and a separate Illumina MiSeq PE 2×250 run (ITS1 region).

#### Abiotic Soil Properties

In May 2020, soil subsamples were stored at 4 °C and shipped to Ward Laboratories within 24 hours of collection for analysis of abiotic soil properties. Specifically, organic matter content was measured using loss on ignition (LOI), soil pH was measured on a 1:1 (w/v) soil to water suspension, and pools of nitrate (NO_3_^-^) were measured by 2M KCl extraction. Cations (K^+^, Ca^2+^, Mg^2+^, and Na^+^) were measured via ammonium acetate extraction and extractable phosphorus (PO_4_^-^) was estimated using the olson method.

### Microbial Bioinformatics

Except where stated otherwise, all DNA sequences were bioinformatically processed using Quantitative Insights into Microbial Ecology 2 (QIIME2; Bolyen et al. 2019). Forward and reverse bacterial/archaeal (16S rRNA) sequences were demultiplexed using the q2-demux plugin then quality filtered, merged, and de-replicated using the q2-dada2 plugin. q2-dada2 also removes PhiX and chimeras before producing sequence clusters with 100% similarity known as amplicon sequence variants (ASV). After quality control, including removal of singletons, taxonomy was assigned to ASVs using the q2-feature-classifier (Bokulich et al. 2018) with the classify-sklearn method against the Silva reference database (Quast et al. 2013).

Fungal (ITS1) sequences were processed using forward reads only, and the ITS1 region was extracted using itsxpress (Rivers et al. 2018) after demultiplexing with the q2-demux plugin. The sequences were then denoised using the q2-dada2 plugin as described above. Singletons were removed along with ASVs that could not be identified to the Kingdom Fungi and those with no BLAST hits. Taxonomy was assigned to ASVs using the q2-feature-classifier with the classify-sklearn method against the UNITE ITS reference database (Nilsson et al. 2018).

### Statistical Analysis

To assess treatment effects on plant biomass, metrics of forage quality, microbial biomass, and abiotic soil properties, we used mean effect sizes and their 95% confidence intervals (CI). An effect size for each paired treated and untreated site within a field was calculated using the natural log of the response ratio:

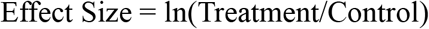

Mean effect sizes and 95% CIs from all 10 fields were then produced by bootstrapping 10,000 replicates in the rcompanion package v2.4.1 (Mangiafico 2017), with the CIs calculated using the basic bootstrap method. This approach offers an intuitive way to interpret treatment effects while providing an alternative to traditional null-hypothesis significance testing (Hubbard and Lindsay 2008; Brennan and Acosta-Martinez 2019). Here, the CIs provide a range of possible effect sizes in which the true treatment effect is likely to reside, and the width of the interval indicates the precision of an estimate. Confidence intervals can also be used to infer significance: when the 95% CIs do not overlap zero, this can be considered as equivalent to a signifiant difference between treated and untreated sites at *α* = 0.05. In some cases, to aid interpretation, we back-transformed the effect sizes and associated CIs into percentage change using the following equation:

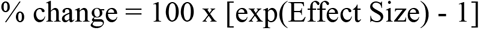

In a single case (i.e., one paired treatment), both the numerator and denominator were 0 and to facilitate inclusion in the effect size calculation, we added a constant of 1 to both.

Treatment effects on bacterial/archaeal and fungal community composition via DNA sequencing were analyzed separate from the other metrics. First, effects were visualized using NMDS of the bray-curtis metric and by plotting grouped pairwise differences in this metric between March 2018 (pre application) and each subsequent sampling date. This visualization exercise was conducted using the vegan package v2.5-7 and ggplot2 package in R (Wickham 2011; Dixon 2003). Treatment effects were then analyzed statistically by using a Kruskal Wallis test to determine whether the pairwise bray-curtis distance between pre and post samples differed by treatment. This analysis was conducted using the longitudinal pairwise-distances command in Qiime2 (Bokulich et al. 2018), which assesses the distance between pre and post sample pairs and determines whether these paired differences differ significantly between two groups.

For a more integrated evaluation of treatment effects, we performed a principal components analysis (PCA) on the full dataset from May 2020 using the prcomp function and factoextra package v1.0.7 in R (Kassambara and Mundt 2020). We paired the PCA with a multi-response permutation procedure (MRPP) based on Euclidean distance to test for a significant treatment difference (Mielke et al. 1981), using the vegan package v2.5-7 (Dixon 2003). The MRPP is a non-parametric procedure that tests the null hypothesis of no difference between groups.

In addition to determining treatment impacts across all 10 fields, for a subset of metrics we also compared effect sizes by field to % SOM values from each control site obtained during the May 2020 sampling event. This approach allowed us to determine whether the response of a given metric to treatment application was moderated by pre-existing levels of this central soil health indicator (Bradford et al. 2019).

## Results

Treatment effects on forage dynamics varied by year and metric. Forage production ranged from 499.2 - 9850.5 kg/ha and was on average 58% higher in treated compared to untreated sites (CI: 25%, 96%; Figure 1). The average difference between treatments remained relatively constant between years (66% and 50% increase with treatment in 2018 and 2019, respectively), but the CI was wider in May 2019 (CI: −0.63%, 117%) than May 2018 (CI: 34%, 104%). In May 2018 and 2019, sites treated with the inoculant and humic product showed elevated levels of forage protein, calcium, and fat content compared to untreated sites—and decreased levels of ADF and NDF (Figure 2). However, these effect sizes were generally small and disappeared in May 2020. In contrast, levels of phosphorus and potassium showed a decline in treated compared to control plots in May 2020, whereas previous to that there were no discernable treatment effects (Figure 2). When averaged across all three years, ADF, calcium, and fat content had means and CIs that did not contain zero. ADF was 4% lower in treated sites (CI: −7.7%, −0.4%), calcium was 18.8% higher (CI: 0.2%, 40.6%), and fat was 6.4% higher (CI: 1.5%, 11.5%).

**Figure 1.**
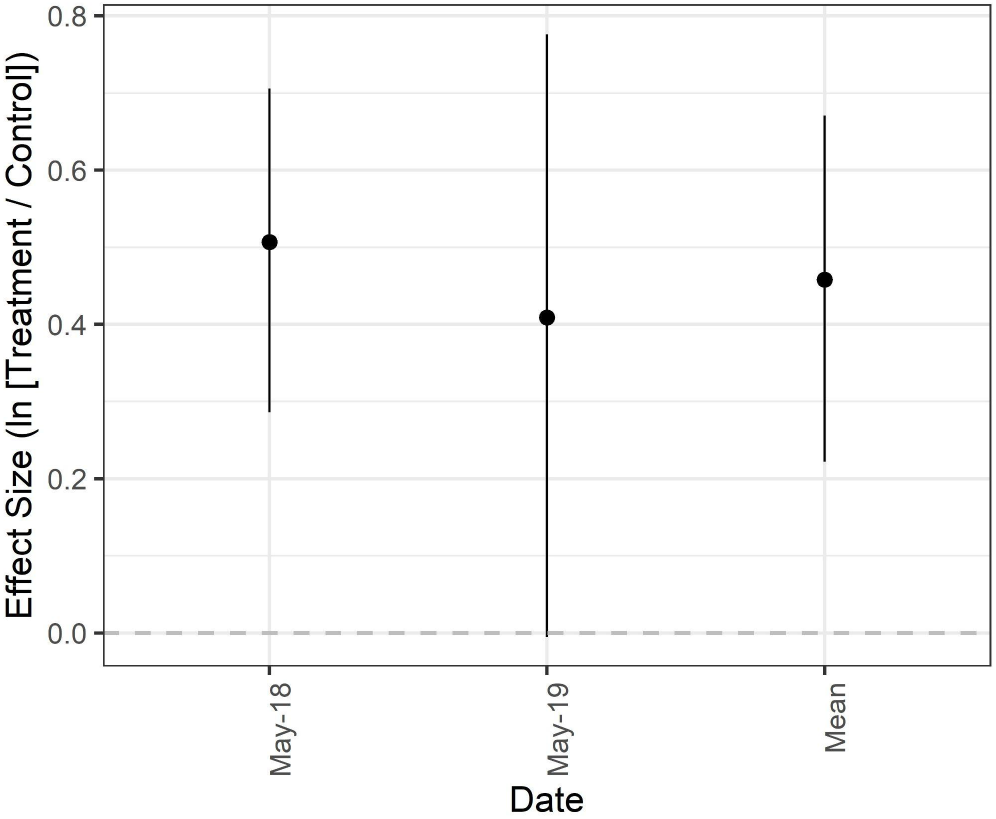
Mean effect size (natural log of the response ratio) and 95% confidence intervals for forage productivity across two years. The mean effect size is a product of all dates combined.

**Figure 2.**
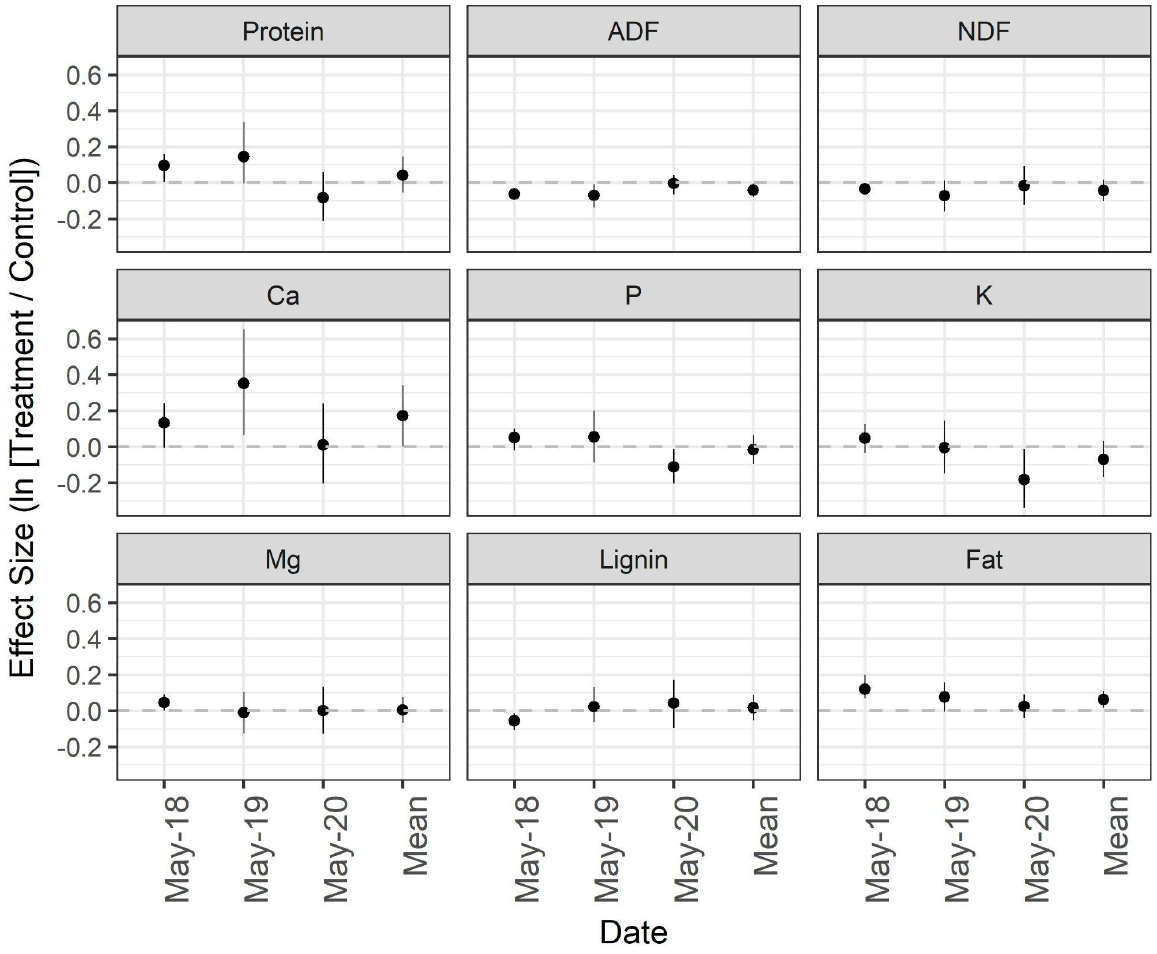
Mean effect size (natural log of the response ratio) and 95% confidence intervals for forage quality metrics across three years. The mean effect size is a product of all dates combined.

Similar to forage, treatment effects on soil microbial communities varied by metric and sampling date. Total fungal biomass ranged from 291.1 - 1599.8 μg/g dry soil and did not show sensitivity to treatment as evidenced by small mean effect sizes and narrow CIs overlapping zero (Figure 3). Active fungal biomass ranged from 1.1 - 265.5 μg/g dry soil and showed the largest difference between treated and untreated sites over time, which can be seen in the comparatively large effect sizes with means ranging from −0.4 to +0.6 across dates. While the data suggest treated sites likely had inherently less active fungal biomass than control sites prior to treatment application, in November 2018 and May 2020, active fungal biomass was on average 40.8% and 86.3% greater in treated compared to control plots (Figure 3). The CI in November 2019 included zero and was relatively wide (CI: −31.1%, 232.0%), but in May 2020 the CI did not include zero (CI: 8.3%, 222.2%), suggesting a significant difference between treatment and control. When averaged across all post-treatment sampling dates, total fungal biomass was 9.1% lower in treated sites (CI: −17.8%, 0.4%) and active fungal biomass was 17.6% higher (CI: −10.1%, 53.57%).

**Figure 3.**
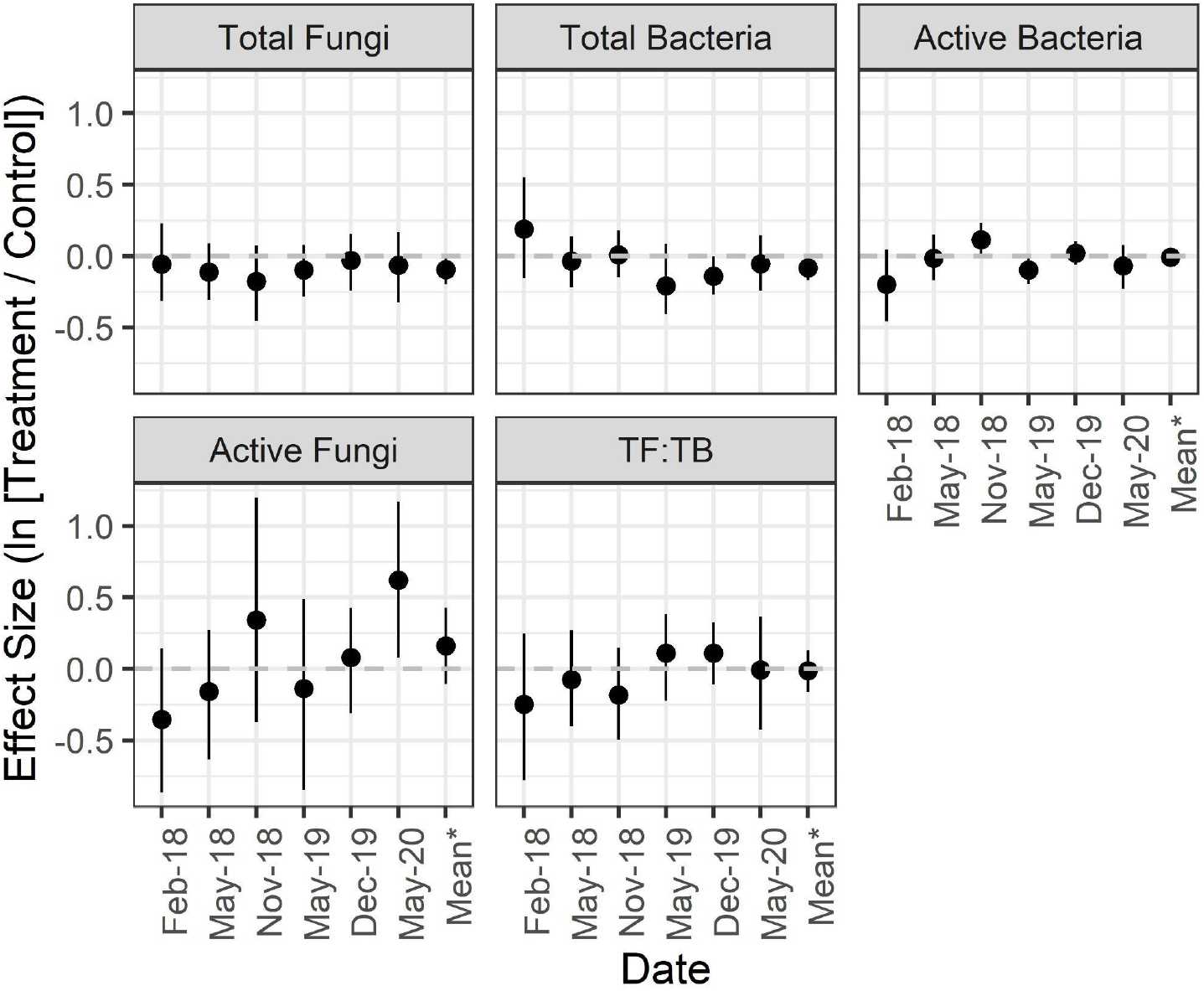
Mean effect size (natural log of the response ratio) and 95% confidence intervals for microbial biomass metrics across three years. *The mean effect size is a product of all dates post-application combined. TF:TB = ratio of total fungi to total bacteria.

Total bacterial biomass ranged from 205.3 - 2893.6 ug/g dry soil and the amount of bacterial biomass in treated compared to control plots decreased slightly after application of the inoculant and humic product. However, effect sizes were small and consistently overlapping zero. Active bacterial biomass ranged from 16.2 - 76.7 ug/g dry soil and the largest difference between sites occurred pre-treatment in February of 2018 (−18.1%; CI: −36.6%, 4.6%). Post-application, the effect sizes for active bacteria oscillated around zero and the CIs of all dates consistently crossed zero with the exception of November 2018, which showed a slight increase in active bacteria in treated compared to control plots (12.2%; CI: 0.11%, 26.2%). The fungi:bacteria ratio varied from 0.2 to 3.8 across all sites and sampling dates, but did not show a strong or consistent response to treatment (Figure 3).

Total protozoa, flagellates, amoeba, and ciliates ranged from 1.2×10^7^ - 6.8×10^8^, 6.3×10^5^ - 5.7×10^7^, 7.1×10^6^ - 6.7×10^8^, and 0 - 1.5×10^6^ individuals/kg dry soil, respectively. Ciliates were the most responsive to application of the inoculant and humic product, with treated sites harboring 131.4% (CI: 18.4%, 343.7%) more ciliates than the control group in May 2020 (Figure 3). For all other protozoan groups and dates, including averages from both dates, the mean effect sizes were in general close to zero with relatively wide error bars that overlap zero, suggesting that treatment had a minimal impact.

To provide a more detailed look at soil microbial community composition, we analyzed DNA amplicon sequences of the 16S rRNA gene and ITS region, which revealed that bacterial/archaeal and fungal community composition remained unaffected by treatment application at this level of resolution. This can be seen visually in the NMDS plots and the pairwise group differences between pre and post treatment application (Figure 6). If treatment-induced differences existed, the distance between pre and post application would be larger for sites treated with the inoculant and humic product than for sites left untreated. The Kruskal Wallis test confirmed that this was not the case and that pairwise bray-curtis distances between pre and post samples did not differ by treatment for bacteria/archaea (Pre-Time1: H = 0.02, P = 0.88; Pre-Time2: H = 0.46, P = 0.49) or fungi (Pre-Time1: H = 0.21, P = 0.65; Pre-Time2: H = 0.09, P = 0.76);

In May 2020, we also assessed abiotic soil properties and ranges of each property can be found in Table 1. Although paired comparisons between the two treatments all included zero, the mean effect sizes and CIs provide evidence that calcium, magensium, and the sum of base cations were more dissimilar between treatments than the other abiotic metrics (Figure 4). For a more holistic and integrated evaluation of treatment effects on abiotic soil properties and other plant and microbial metrics collected in May 2020—the final sampling date in our study—we paired a PCA with a MRPP test of significance. The MRPP illustrated that, when all metrics were taken into account, treated and control sites did not differ significantly (effect size *A* = 0.03; P = 0.18). This also can be observed visually with the PCA, which shows considerable overlap between the two groups (Figure 5).

**Table 1.** Abiotic soil properties (mean, standard deviation, min, and max values)

| Metric | Average | Standard Deviation | Min | Max |
| --- | --- | --- | --- | --- |
| Soil pH | 5.18 | 0.29 | 5.20 | 6.4 |
| SOM | 6.14 | 1.46 | 3.60 | 9.3 |
| Nitrate | 2.73 | 2.11 | 0.40 | 8.2 |
| Phosphorus | 16.20 | 17.35 | 4.70 | 67.7 |
| Potassium | 343.55 | 86.55 | 209.0 | 489.0 |
| Calcium | 2051.7 | 716.57 | 712.0 | 3268.0 |
| Magnesium | 758.05 | 283.52 | 358.0 | 1453.0 |
| Sum of Cations | 26.36 | 4.13 | 17.30 | 34.6 |

**Figure 4.**
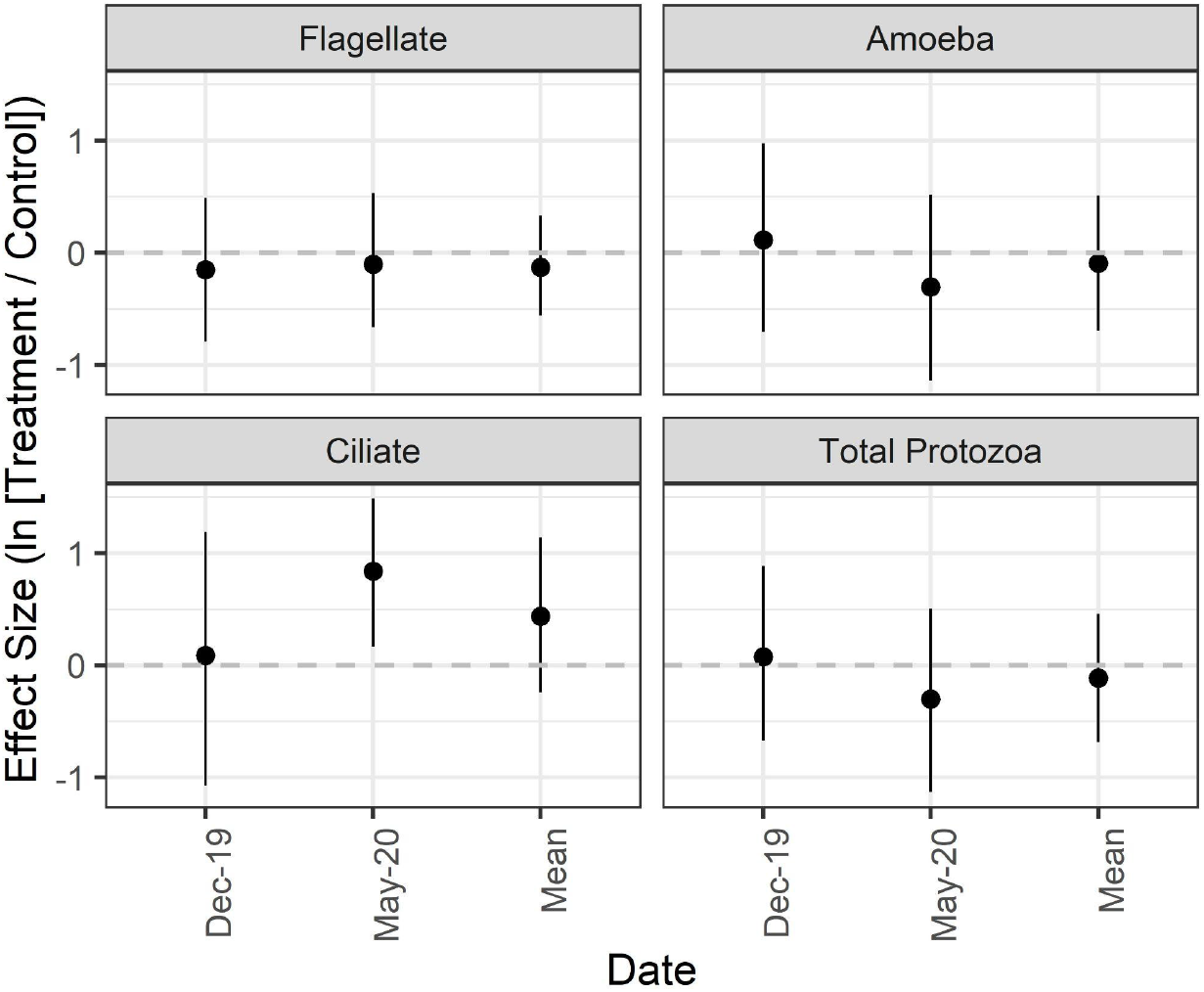
Mean effect size (natural log of the response ratio) and 95% confidence intervals for protozoa across three years. *The mean effect size is a product of all dates post-application combined.

**Figure 5.**
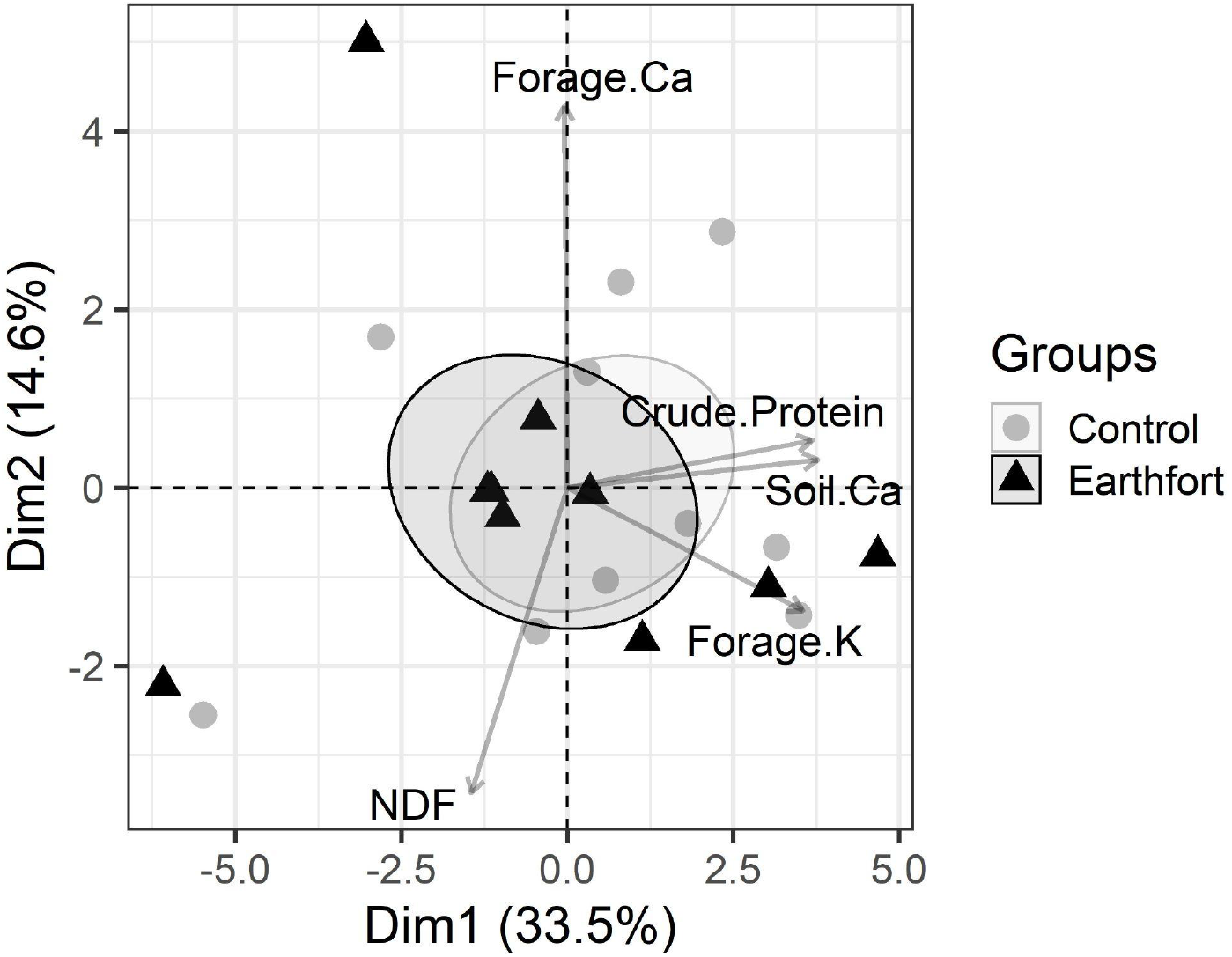
Principal components analysis (PCA) of plant, microbial, and soil metrics collected in May 2020. Symbols denote treatment (Black triangle = treatment with the Earthfort inoculant and humic product; Gray circle = untreated control) and ellipses correspond to 95% confidence intervals. Vectors denote metrics, of which only the top 5 contributing ones are listed.

**Figure 6.**
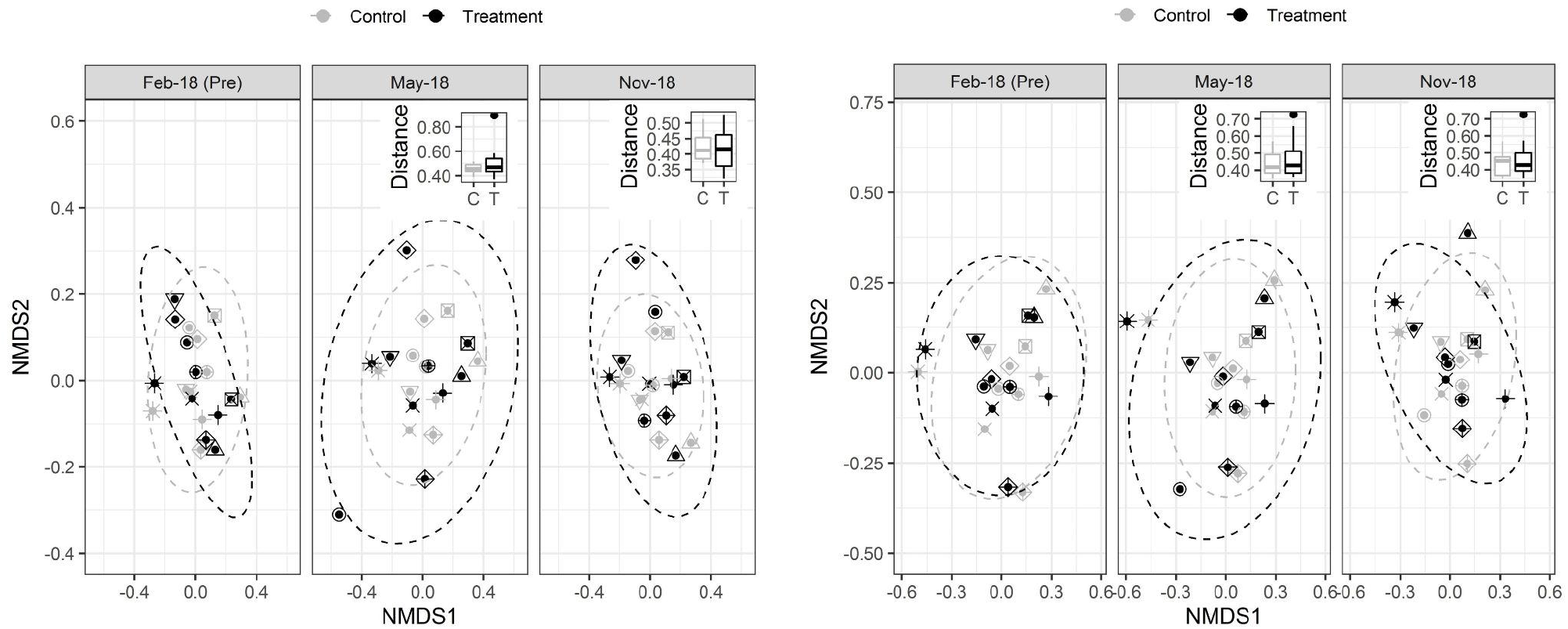
PCoA plots of bacteria/archaea (left panel) and fungi (right panel) with pairwise distance comparisons as insets.

## Discussion

Understanding how commercially available biostimulant products, such as microbial inoculants and humic substances, influence above and belowground dynamics on rangelands is important, especially given increasing interest in these products as a way to help manage rangelands regeneratively. In this study, we examined the combined effects of two such products that are currently on the market, and asked whether they influenced rangeland forage productivity and quality, soil microbial biomass and community composition, and abiotic soil parameters in Central Coastal California. While field studies such as this are limited in their ability to disentangle the chemical and biological mechanisms underpinning observed effects, they are useful for evaluating products under real-world conditions, assessing trade-offs, and generating a better understanding of the potential to reap economic and ecological benefits across varying agricultural production systems.

We found that foliar application of the commercial inoculant and humic product consistently enhanced within-season forage production. Given only three out of ten fields received microbial inoculant as part of this first amendment, it is likely that the humic substance drove the observed response. However, it is possible that introduction of the endophytic bacteria *Bacillus subtilis* or other microbial taxa in the inoculant contributed to plant growth promotion as well, particularly in the second year when all sites received both products (Preininger et al. 2018). Very few studies have measured how forage productivity changes in response to humic substances (Verlinden et al. 2007), and of those, most isolate one focal plant species (Churkova 2013; Khaleda et al. 2017). Verlinden et al. (2010) found that applying a mixture of humic and fulvic acids increased initial grass production at four pastures dominated by *Lolium multiflorum* and *L. perenne* in northern Belgium; however, this effect attenuated after the first cut. The size of the initial response (i.e., the natural log of the response ratio) reported by Verlinden et al. (2010) ranged from 0.07 - 0.14, depending on the mode of application, which is considerably less than the mean effect sizes 0.51 in 2018 and 0.41 in 2019 from our study. Our work joins this small but growing list of others to illustrate that there is a potential role for humic products to play when seeking to boost forage productivity of pasture and rangeland systems.

Assessing effect sizes is a critical aspect of interpreting the ecological and economic significance of treatment application for any outcome (e.g., Rinella and James 2017). This is particularly true on agricultural lands, where decisions about how to allocate limited time and funding to achieve regenerative outcomes must be made carefully. When transformed to percentage change, forage production in treated sites was on average 65% (CI: 34%, 104%) and 50% (CI: −0.6%, 117%) higher than control sites in 2018 and 2019, respectively. This is equivalent to an average increase in forage production of 1659 kg/ha and 1314 kg/ha. This level of change between treatment and control is equivalent to what can be seen across growing seasons with different amounts of rainfall on the Central Coast (Becchetti et al. 2016). This shows the potential for the product to boost forage production, with possible benefits for livestock stocking density and carrying capacity. While the results need to be confirmed in other contexts before recommendation at scale, the promising nature of them warrants increased research and consideration of deploying this product in an adaptive management framework to help manage forage production on rangelands.

We found no evidence that background levels of SOM influenced how forage productivity responded to treatment application, despite a more than two-fold difference in SOM levels across sites. Soil organic matter provides myriad benefits including improved water retention and nutrient supply to plants (reviewed in Lal 2009) as well as support of soil biodiversity (Murphy et al. 2011). Soils with higher SOM levels should therefore create conditions more conducive to plant growth and resiliency, all else equal. Oldfield et al. (2019) demonstrated this by comparing maize and wheat yields to soil organic carbon (SOC; a proxy for SOM) globally and finding that yields correlated positively with SOC until a threshold of 2% (Oldfield et al. 2019). A recent study by Kane et al. (2021) demonstrated that the benefits of SOM—namely benefits to available water capacity and cation exchange capacity—extend beyond this to help stabilize maize yields during stressful drought conditions. Because less optimal growing conditions are likely to enhance the benefits of microbial inoculation (Hart et al. 2018) and humic substances (Olk et al. 2021), it is conceivable that sites with lower SOM levels would be in greater need and thus more responsive to application. However, we found no evidence for this to be the case, providing new information that at least in rangelands of Central Coastal California, this contextual parameter does not seem to moderate the impact of treatment application on peak growth. Other aspects of forage production, such as growing season length, are important in these systems from a livestock production standpoint, and it’s possible that product application could interact with contextual factors like SOM to influence this and other parameters not captured here.

In addition to boosting forage productivity, the inoculant and humic product improved some aspects related to forage quality—albeit to a lesser degree. Notably, concentrations of ADF were 4.0% lower in treated compared to untreated sites when averaged across all three years. Acid detergent fiber contains cellulose and lignin, and is inversely related to the digestibility and energy of forage (Colburn and Evans 1967). While an increase in crude fiber has been observed in crops like corn and soybean (Kocira et al. 2019; Efthimiadou et al. 2020), a decrease in ADF with application of the inoculant and humic product in this case suggests an improvement to forage quality for rangeland systems. Further confirming this, calcium was on average 18.8% higher and fat content was on average 6.4% higher in sites that were treated compared to those that were not. Fat is a potent source of energy and calcium serves as an essential plant nutrient, and increases in these metrics can be interpreted as indicating improved forage quality. Indeed, although beef cattle in California tend to be deficient in other minerals like manganese, zinc, and selenium (Davy et al. 2019), deficiencies in calcium can affect the livestock industry and so increased concentrations of this nutrient may be important in some cases. While we cannot disentangle the mechanisms behind such a response, it’s possible that the humic product stimulated uptake of calcium (Olk et al. 2018), which is generally immobile in plants, and influenced metabolic pathways that regulate lipid, lignin, and cellulose production. Regardless of the mechanism(s), these findings suggest that application of a microbial inoculant and humic product combination have the potential to improve some components of forage quality and palatability in rangeland systems.

Unlike aboveground forage dynamics, we did not see a strong or consistent response to treatment application for soil microbial communities or abiotic soil parameters. Given that the microbial inoculant was applied as a foliar spray, it is perhaps not surprising that belowground shifts in microbial biomass, activity, or community composition were not apparent within the timeframe of our study. Microorganisms applied in this way are thought to improve plant growth, quality, and stress resilience by entering the plant via the stomata (Preininger et al. 2018). They are not likely to intercept and sufficiently interact with the soil when applied at low concentrations to sites that have considerable live or dead standing plant material, such as those included in this study. Similarly, because we were applying humic products as a biostimulant, the application rates were too low to influence SOM dynamics or nutrient availability via direct addition. However, as others have suggested (Olk et al. 2018), there is a potential for soil health metrics like microbial biomass and SOM to improve over time if plant productivity remains elevated, especially of root growth (Conselvan et al. 2017; Rasse et al. 2005). Given the importance of soils for sustaining ranch operations and public benefits like climate change mitigation (Bradford et al. 2019), the direct and indirect benefits of these management tools will be important to capture in the near- and long-term to inform expectations and decision-making for ranchers and policy-makers who are looking for new products to help build resiliency into landscapes.

## Conclusion

There is increasing interest in using biostimulant products, such as microbial inoculants and humic substances, to help manage rangelands regeneratively. Understanding how above and belowground dynamics on rangelands respond to these products is therefore important. In our study, we found that forage productivity and some metrics of forage quality responded positively to the foliar application of a commercial microbial inoculant and humic product. These benefits were not mirrored by changes belowground in the microbial community or abiotic parameters, but also did not come at a cost to the measured parameters as well. Depending on the goals of using the products, this could be seen as a winning scenario and suggests microbial inoculants and humic products could warrant attention as a potential tool for regenerative stewardship of rangelands. While our study derives from one ranch and therefore requires confirmation of its ubiquity, our results provide compelling new insights into the usefulness of this approach for managing rangeland productivity in California’s Central Coast. As with any new management tool, we recommend ranchers interested in these products first apply to small areas in an adaptive management framework and then scale accordingly.

## Acknowledgements

Acknowledge field help and reviewers of manuscript. We thank many TomKat apprentices including Andrea Hatsukami, Justin Miller, Drake Swezey, Evan Watson, Dillon Gruber, Jessica Teresi, Alex Michel, Jose Ramirez Martinez, Mariana Zavala, Lana Jansen, and Celia Hoffman. We also thank Wendy Millet, Libby Porzig, Grant Ballard, Suzane Rowland for support of the project and review of the manuscript.

