## Appendix for "Applying biostimulants boosts forage productivity without affecting soil biotic and abiotic parameters on a Central Coast California rangeland"

Appendix Figure 1. Annual cumulative rainfall derived from a weather station at TomKat Ranch for each year of the study. Water years were defined as July 1-June 30.

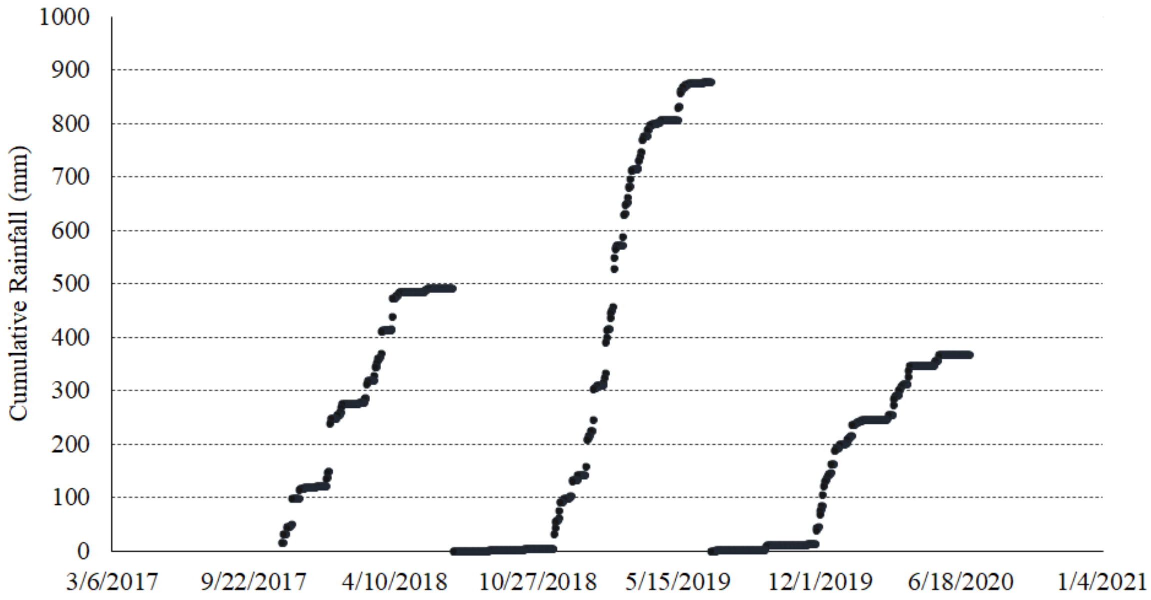


Appendix Figure 2. Experimental design schematic. Paired treated and control areas were 1 acre each and replicated across ten fields. Soil samples were collected from within 10m x 10m areas near the center of each acre. Grazing exclosures were located adjacent to the center square and rotated to a new area each year.

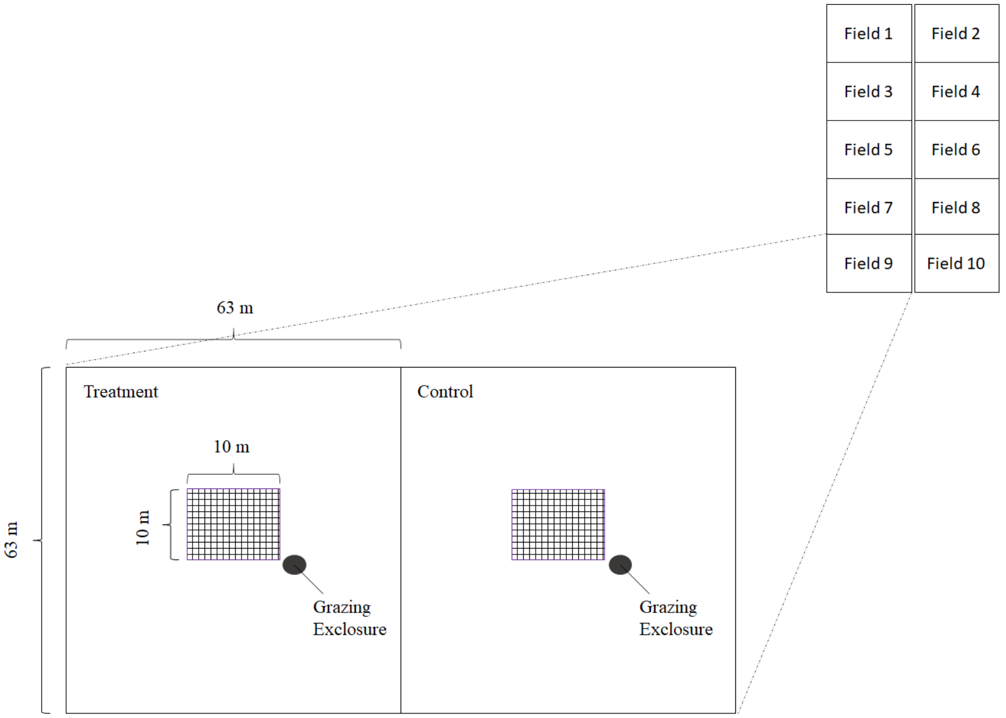

Appendix Figure 3. Timeline of treatment application and plant and soil sampling. Brown = only soil sampling. Green = plant and soil sampling.

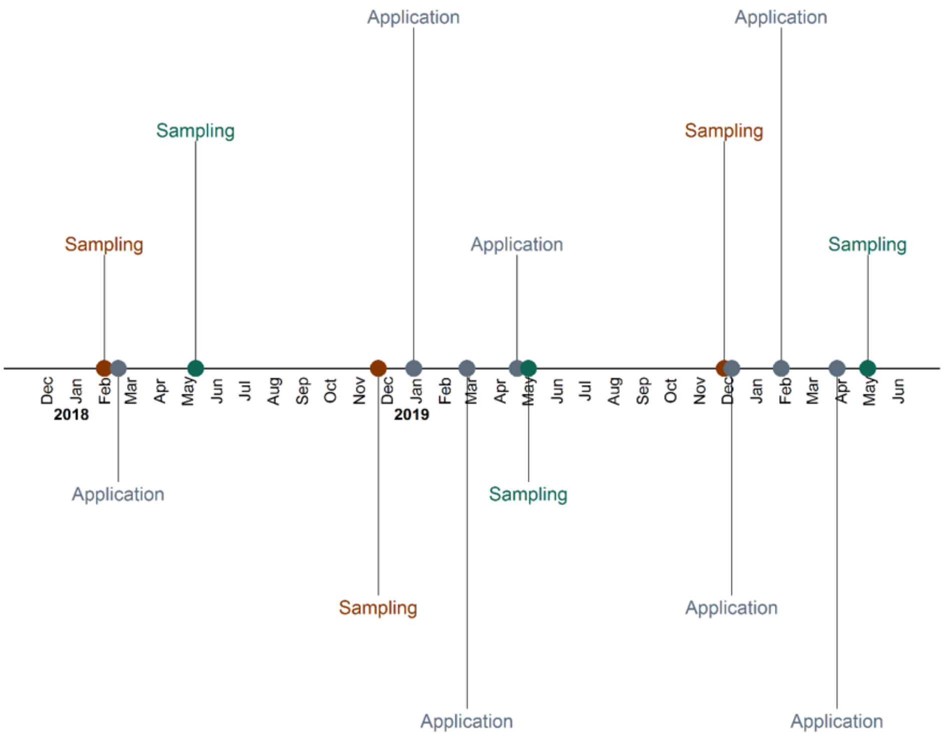


Appendix Figure 4. Relationship between treatment effects on forage productivity and % SOM by field.

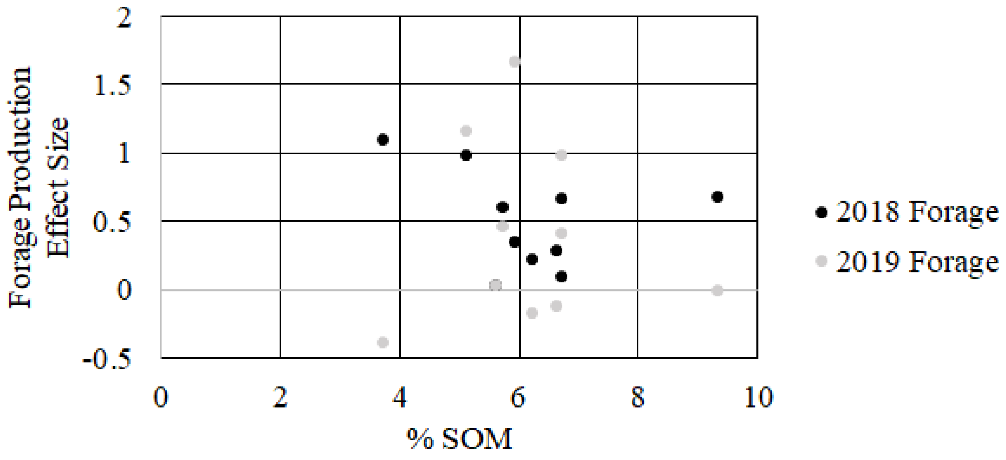

Appendix Table 1. Rates for first application between February 20-24, 2018. During subsequent applications, all fields received a constant 1.12 kg provide and 9.35 liters revive in 280 liters water per hectare.

| | **Application Rates** | |
|---|---|---|
| **Field** | **Provide (Inoculant) kg/ha** | **Revive (Humic) liters/ha** |
| Fertility Flat | 0 | 9.35 |
| Front Field | 0 | 9.35 |
| Lane Hill | 1.12 | 9.35 |
| Lone Tree | 1.12 | 4.69 |
| Mike's Meadow | 0 | 9.35 |
| Moore | 0 | 4.69 |
| PRBO | 0 | 9.35 |
| Stage South | 1.12 | 4.69 |
| Upper China | 0 | 9.35 |
| Water Tank | 0 | 4.69 |
